# FAM13A regulates KLRG1 expression and interferon gamma production of natural killer cells

**DOI:** 10.1101/2020.08.26.268490

**Authors:** Ni Zeng, Maud Theresine, Christophe Capelle, Neha D. Patil, Cécile Masquelier, Caroline Davril, Alexandre Baron, Djalil Coowar, Xavier Dervillez, Aurélie Poli, Cathy Leonard, Rudi Balling, Markus Ollert, Jacques Zimmer, Feng Q. Hefeng

**Author notes:** Correspondence should be addressed to J.Z. or F.Q.H.

## Abstract

The polymorphism of the gene *FAM13A* (family with sequence similarity 13, member A) is strongly linked to the risk of lung cancer and chronic obstructive pulmonary disease, which are among the leading causes of mortality and morbidity in lung-related diseases worldwide. However, the underlying molecular and cellular mechanisms through which *FAM13A* contributes to the pathogenesis of these diseases largely remain unclear. Here, using a *Fam13a* knock out (KO) mouse model, we showed that *Fam13a* depletion upregulated the expression of the terminal differentiation and inhibitory marker, KLRG1 (killer cell lectin-like receptor G1) in natural killer (NK) cells. NK cells from *Fam13a*-deficient mice showed impaired IFN-γ production either against target tumor cells or following various cytokine cocktail stimulations. Furthermore, the number of lung metastases induced by B16F10 melanoma cells was increased in *Fam13a*-KO mice. Collectively, our data suggest a key role of *FAM13A* in regulating NK cell functions, indicating that the key lung-disease risk gene *FAM13A* might contribute to the pathogenesis of several lung diseases via regulating NK cells.

## Introduction

Several independent genome-wide association studies (GWAS) among various populations have already reproducibly shown the strong link between the polymorphism of *Family with sequence similarity 13, member A (FAM13A)* and chronic obstructive pulmonary disease (COPD) and lung cancer, the leading entities causing mortality in lung-related diseases worldwide [1–12]. Functional studies have revealed that *Fam13a* depletion reduces the susceptibility to COPD via inhibiting the WNT/β-catenin pathway in a mouse model [13]. In human lung tumor cell lines, knocking-down *FAM13A* reduced tumor cell proliferation but induced cell migration *in vitro* [14]. However, the exact *in vivo* physiological role of FAM13A in the complicated process of tumor onset and metastasis still remains mysterious. Recently, a few studies indicated that FAM13A might be involved in the regulation of immune responses. Eisenhut et al. observed the upregulation of FAM13A in CD4^+^CD25^−^ effector T cells but reduced expression in T regulatory cells (Tregs) of human blood [14]. In contrast, in our previous study [15], the expression of FAM13A was increased in human Tregs vs. CD4^+^ effector T cells following TCR stimulation. Another work using the expression quantitative trait loci (eQTL) method has briefly investigated the functional effect of *FAM13A* knockdown on human naïve CD4^+^ T cells *in vitro* [16]. Meanwhile, microRNA-328 in M2 macrophage-derived exosomes has been demonstrated to regulate the progression of pulmonary fibrosis via *Fam13a* in an animal model [17]. Those different reports all suggest potential roles of *Fam13a*, although sometimes even controversial, in immune cells, which, however, highlights a need of more investigation to clarify further the complicated *invivo* functions of *Fam13a* in the immune system.

Here we utilized a *Fam13a* whole-body knockout (KO) mouse model to study the potential *in vivo* impact of *Fam13a* on cellular phenotypes and functions of major immune cells, including NK cells, T and B lymphocytes. We found that *Fam13a* did not affect the homeostatic composition and basic functional markers of total B cells, CD4^+^ T cells, CD4^+^ Tregs and CD8^+^ T cells. Interestingly, *Fam13a* depletion upregulated the expression of the critical maturation and inhibitory marker, KLRG1 [18, 19] while impairing the IFN-γ production of NK cells. Notably, *Fam13a* depletion exacerbated lung metastasis induced by B16F10 melanoma cells in the C57BL/6 (B6) mouse model. Altogether, our results provide strong evidence that *Fam13a* is an important component regulating the effector functions of NK cells, mostly through modulating KLRG1 expression and IFN-γ production.

## Results

### *Fam13a* depletion has no spontaneous effects on homeostatic cellularity and functions of major immune subsets

To identify the potential effect of *Fam13a* on the homeostatic phenotypes of the immune system, we first analysed the composition of different major immune cell types in various lymphoid organs and tissues of *Fam13a* KO mice [Fam13a^tm2a (KOMP)Wtsi^] with B6 background obtained from the Knockout Mouse Project. The same mice have been characterized elsewhere [13], but not in the context of immunology. As a starting point, we compared the mRNA expression of *Fam13a* in the lung tissue between *Fam13a* knockout (KO) and wild type (WT) mice and that in NK cells isolated from spleen. As expected, *Fam13a* KO mice relative to WT littermates exhibited a clear reduction in the transcript expression of *Fam13a* in both the lung tissue (**S1a Fig**) and NK cells (**S1b Fig**). We checked the frequency of CD19^−^CD3^−^NK1.1^+^ NK cells in several relevant tissues including spleen (**S1c and 1d Fig**), peripheral lymph nodes (pLNs) (**S1e Fig**), bone marrow (BM) (**S1f Fig**) and lung (**S1g Fig**) and no significant difference between homeostatic *Fam13a* KO and WT mice was observed. We also could not see any significant difference in the frequencies of several other major types of lymphocytes, such as CD3^+^ total T cells, CD19^+^ B cells, CD8^+^ T cells, CD4^+^ T cells as well as CD4^+^ Treg cells (**S1 Table**). We further characterized the naïve and memory compartment of CD4^+^ and CD8^+^ T cells. Again, no significant difference was found in the percentages of naïve (CD44^low^CD62L^high^) and effector memory (CD44^high^CD62L^low^, EM) CD4^+^ T cells and naïve, EM and central memory (CD44^high^CD62L^high^, CM) CD8^+^ T cells at the tested age (8-12 wks) (**S1 Table**). In conclusion, *Fam13a* depletion does not lead to spontaneous abnormalities in the development of various major immune cells under homeostasis.

To further evaluate whether *Fam13a* influences the basic homeostatic functions of those immune cells, we analysed the expression of an activation marker (CD69), an exhaustion/activation marker (PD-1) and a proliferation marker (Ki-67) of CD4^+^ and CD8^+^ T cells. No significant difference in the frequency of the cells expressing any of those markers among CD4^+^ T cells and CD8^+^ T cells in spleen exhibited between *Fam13a* KO and WT mice (**S1 Table**). We also analyzed a key subset of T cells, Tregs, which play a key role in suppressing responses of effector immune cells. Therefore, we evaluated their suppressive function by co-culturing the CFSE-labelled CD4^+^ conventional T cells (Tconv), antigen-presenting cells (APCs) and Tregs. Loss of *Fam13a* did not compromise Treg suppressor function against Tconv proliferation (**S2a Fig**). In short, *Fam13a* depletion causes deficiency neither in the development, nor in the activation or proliferation of major lymphoid subsets such as CD4^+^ and CD8^+^ T cells, at least in the analyzed mice lines aged 8-12 weeks under homeostatic conditions.

### *Fam13a* depletion upregulates the expression of KLRG1 in NK cells

As demonstrated above, *Fam13a* depletion did not generate differences in the frequency of total NK cells under homeostatic conditions, which, however, cannot exclude the potential effect of *Fam13a* on certain NK functions. To explore whether *Fam13a* affects major functional markers of NK cells, we first investigated the maturation profile of NK cells by checking the co-expression of CD27 and CD11b. Interestingly, we noticed that *Fam13a* depletion led to a significant but only modest change in the proportions of immature CD11b^−^CD27^+^and mature CD11b^+^CD27^−^ NK cells (**S3a-c Fig**). Since CD11b^+^CD27^−^ NK cells are mature NK cells, representing the major KLRG1-expressing NK subset [19–21], we also analyzed the expression of the inhibitory receptor KLRG1 among NK cells. In line with the data related to CD27 and CD11b subsets, we observed a much higher percentage of KLRG1-expressing cells among total NK cells in both spleen (**Fig 1a, b**) and pLNs (**Fig 1c, d**) of the *Fam13a*-KO mice.

**Fig 1.**
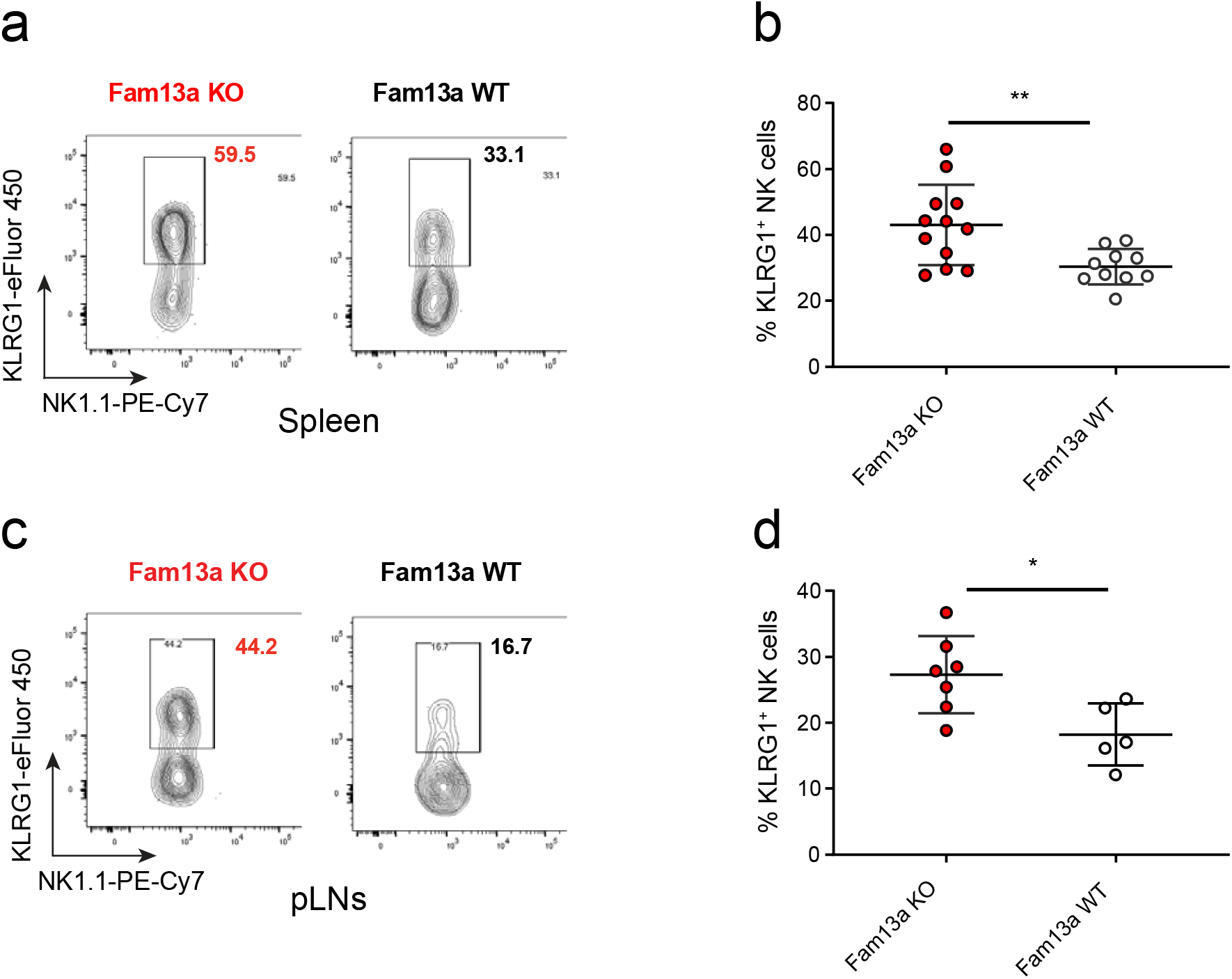
*Fam13a* depletion upregulates the expression of KLRG1 in NK cells. **(a)** Representative FACS plots of KLRG1 and NK1.1 expression on CD19^−^CD3^−^ NK1.1^+^cells in spleen. **(b)** Percentages of KLRG1^+^CD19^−^CD3^−^ NK1.1^+^ among NK cells in spleen of *Fam13a* KO and WT littermates (KO, n=12; WT, n=10). **(c)** Representative FACS plots of KLRG1 and NK1.1 expression on CD19^−^CD3^−^ NK1.1^+^cells in pLNs of *Fam13a* KO and WT littermates. **(d)** Percentage of KLRG1^+^ subpopulation among NK cell in pLNs of *Fam13a* KO and WT littermates (KO, n=7; WT, n=5). Results represent four independent experiments. Data are mean ± s.d. The p-values were determined by a two-tailed Student’s t-test. n.s. or unlabeled, not significant, *p<=0.05, **p<=0.01 and ***p<=0.001.

To get a more comprehensive picture, we further analyzed different inhibitory receptors (IR) and activation receptors (AR) of NK cells, whose engagement is critical to regulate and balance NK cell activities. For the AR, the frequency of NKp46^+^, Ly49H^+^ and Ly49D^+^ NK cells (**S3d-f Fig**), as well as the expression of 2B4 and NKG2D (**S3g-h Fig**) were not significantly altered in *Fam13a* KO NK cells. For the IR, we did not observe any significant difference in the expression of Ly49A, Ly49C/I and NKG2A (**S3i-k Fig**) on NK cells between *Fam13a* KO and WT mice. In summary, loss of *Fam13a* critically upregulated KLRG1 expression, indicating that *Fam13a* has an inhibiting or regulatory effect on NK cell maturation or preventing to some extent premature entry into the senescent state.

### *Fam13a* depletion impairs NK-cell IFN-γ production

A few studies have already demonstrated that KLRG1 inhibited NK-cell IFN-γ production as well as cytotoxicity [18, 22, 23]. Since we observed a significant upregulation in the frequency of the KLRG1^+^ NK cells, we further assessed IFN-γ production and degranulation of NK cells. We expanded NK cells by culturing total splenocytes isolated from *Fam13a* KO or WT littermates in the presence of a high concentration of IL-2 for 5 days. Then the cells were restimulated via different cytokine cocktails to check IFN-γ production **(Fig 2a)**. We found that *Fam13a* deficiency did not affect the survival and expansion of NK cells *in vitro* (**Fig 2b-c**). However, when stimulated with the cytokine cocktail containing IL-2, IL-12 and IL-15, *Fam13a*-deficient NK cells produced a significantly lower amount of IFN-γ compared with their WT counterparts (**Fig 2d-e**). Furthermore, even when stimulated with the strong activation cytokine cocktail composed of IL-2, IL-12 and IL-18, the deficiency in IFN-γ production capacity *of Fam13a*-KO NK cells was not compensated (**Fig 2f-g**). Taken together, our data demonstrate that *Fam13a* depletion impairs IFN-γ production in NK cells under various cytokine-cocktail stimulations *in vitro*.

**Fig 2.**
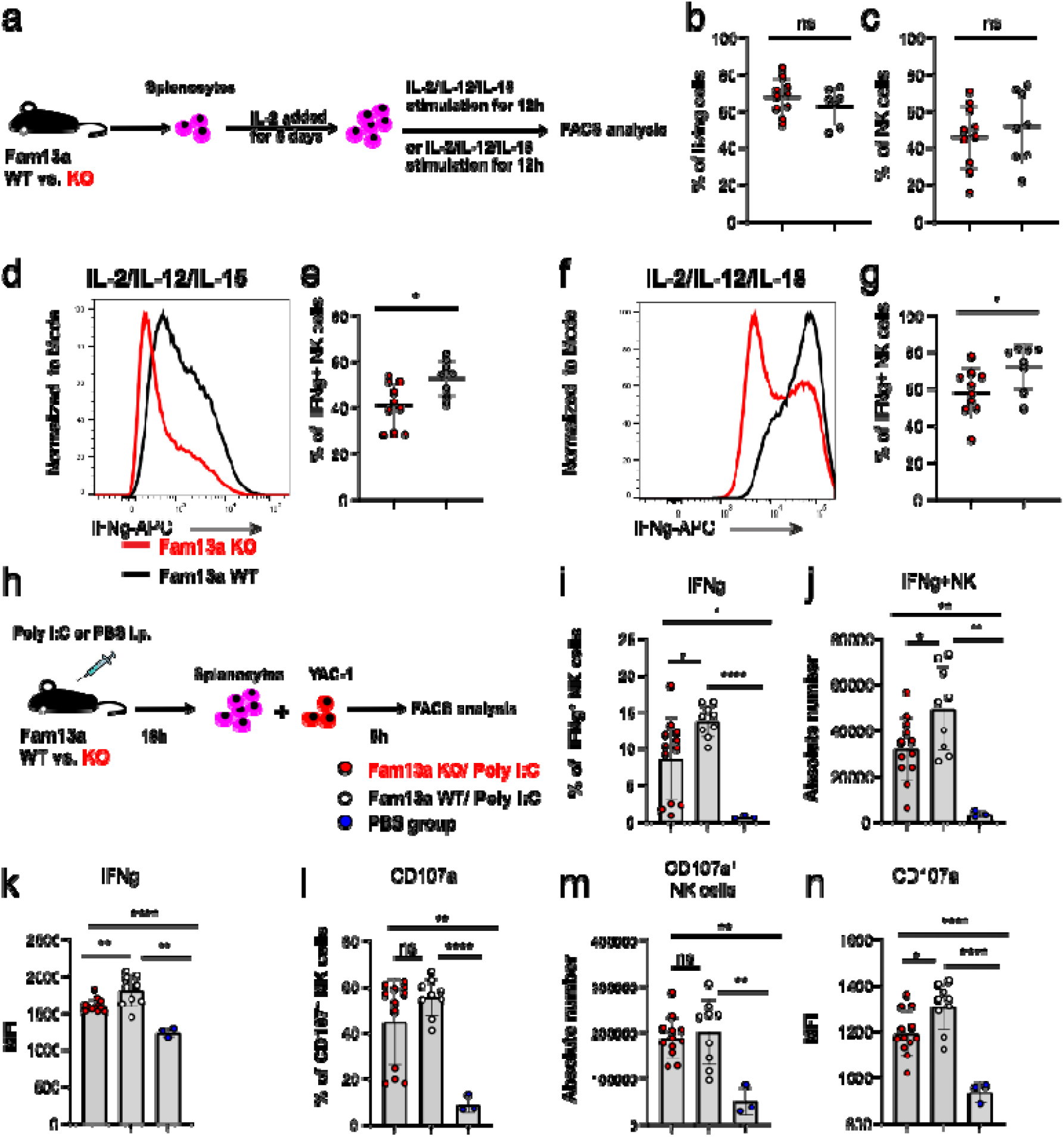
*Fam13a* deficiency impairs NK-cell IFN-γ production following *in vitro* or *ex vivo* stimulation. **(a)** Schematic of the experimental setup to stimulate NK cells with different cytokine cocktails. **(b)** Percentage of living cells after 5-day expansion in the presence of IL-2 (KO, n=10; WT, n=7). **(c)** Percentage of NK1.1^+^ NK cells after 5-day expansion in the presence of IL-2 (KO, n=10; WT, n=8). **(d)** Representative FACS histogram overlay of IFN-γ production of *Fam13a* KO (red line) and WT (black line) NK cells following 5-day IL-2 expansion and IL-2, IL-12 and IL-15 re-stimulation. **(e)** Percentage of IFN-γ producing cells among total NK cells after IL-2, IL-12 and IL-15 re-stimulation (KO, n=12; WT, n=9). **(f)** Representative FACS histogram overlay of IFN-γ production of *Fam13a* KO (red line) and WT (black line) NK cells after 5-day IL-2 expansion and IL-2, IL-12 and IL-18 re-stimulation. **(g)** Percentage of IFN-γ producing cells among total NK cells after IL-2, IL-12 and IL-18 re-stimulation. **(h)** Schematic of the experimental setup to analyze the ex-vivo degranulation capacity of NK cells. *Fam13a* KO or WT littermates were injected intraperitoneally (i.p.) with 150 µg poly(I:C) or PBS and then the next day splenocytes were incubated with YAC-1 cells to evaluate NK cell IFN-γ production and degranulation. **(i)** Percentage of IFN-γ producing NK cells of *Fam13a* KO and WT NK cells against YAC-1 tumor cells. **(j)** Absolute number of IFN-γ producing NK cells of *Fam13a* KO and WT NK cells against YAC-1 tumor cells. **(k)** Geometric mean of IFN-γ expression intensity in total IFN-γ^+^ NK cells of *Fam13a* KO and WT NK cells. **(l)** Percentage of CD107a producing NK cells of *Fam13a* KO and WT NK cells against YAC-1 tumor cells. **(m)** Absolute number of CD107a producing NK cells of *Fam13a* KO and WT NK cells against YAC-1 tumor cells. **(n)** Geometric mean (MFI) of CD107a expression intensity in total CD107a^+^ NK cells of *Fam13a* KO and WT NK cells (KO, n=12, WT, n=9). Results are representative of three (**b, c, e, g**) and two (**i-n**) independent experiments. Data are mean ± s.d. The *p*-values were calculated by a two-tailed Student’s t-test. n.s. or unlabeled, not significant, \**p*<=0.05, \*\**p* <=0.01 and \*\*\**p* <=0.001.

To further check the killing capacity of NK cells towards tumor cells, we pre-activated NK cells *in vivo* by injecting the TLR3 agonist poly(I:C) into *Fam13a* KO and WT littermates (**Fig 2h**). We then evaluated the expression of IFN-γ and the degranulation marker CD107a of NK cells following the incubation with the target tumor cell line YAC-1. In line with the *in vitro* observations, lower amounts of IFN-γ were produced by *Fam13a* KO NK cells, as shown by different readouts, such as the lower frequency of IFN-γ-expressing NK cells (**Fig 2i**), the smaller absolute number of IFN-γ-expressing NK cells (**Fig 2j**) and the decreased MFI of IFN-γ per NK cell in IFN-γ^+^ NK cells (**Fig 2k**). CD107a reflects the cytotoxic activity of NK cells and cytotoxic CD8^+^ T lymphocytes [24, 25]. No significant difference was observed in the percentages of CD107a-expressing cells among total NK cells (**Fig 2l**) and in the absolute number of CD107a^+^ NK cells (**Fig 2m**) between *Fam13a* KO and WT mice following poly(I:C) injection. Interestingly, loss of *Fam13a* caused a modest but significant decrease in the degranulation capacity per individual NK cell against YAC-1, as indicated by the decreased MFI of CD107a in *Fam13a* deficient NK cells (**Fig 2n**). In conclusion, *Fam13a* depletion essentially impairs NK-cell IFN-γ production against YAC-1 tumor cells.

### *Fam13a* deficiency worsens lung metastasis induced by melanoma cells *in vivo*

Having shown that *Fam13a* depletion impaired certain critical effector functions of NK cells *in vitro* and *ex vivo*, we sought to further investigate the *in vivo* effects. Since NK cells are critical in the control of tumor metastasis [26], we induced lung metastasis by intravenous injection of B16F10 melanoma cells into *Fam13a* whole-body KO and WT littermates. After 16 days post inoculation, we evaluated the lung metastases (**Fig 3a**). Compared with WT littermates, *Fam13a* KO mice had developed much more tumor metastases in the lung (**Fig 3b-c**). For cellular composition phenotypes, we first analyzed NK cells in the spleen and lung tissues. The frequency and absolute number of NKp46^+^NK1.1^+^ NK cells among CD3^−^CD19^−^ splenocytes was much lower in B16F10-inoculated *Fam13a* KO mice compared with that in WT littermates (**Fig 3d-f**). Importantly, *Fam13a* KO mice also had a lower percentage of infiltrated NKp46^+^NK1.1^+^ NK cells among CD3^−^CD19^−^ cells in the tumor tissue, i.e., lung in comparison to WT littermates (**Fig 3g**). Furthermore, similar to the homeostatic phenotype of *Fam13a* KO NK cells (**Fig 1a-b**), a much higher frequency of KLRG1^+^ cells were also observed among total NK cells in both spleen (**Fig 3h-i**) and lung (**Fig 3j**) of *Fam13a* KO mice in comparison with that in WT littermates. In the chronic infection model, such as chronic hepatitis C virus infection, KLRG1^+^ NK cells represent an exhausted phenotype and lower capacity for IFN-γ production [27–29]. This indicates that both the decreased number and the high level of KLRG1 of NK cells in *Fam13a* KO mice together aggravate tumor metastasis.

**Fig 3.**
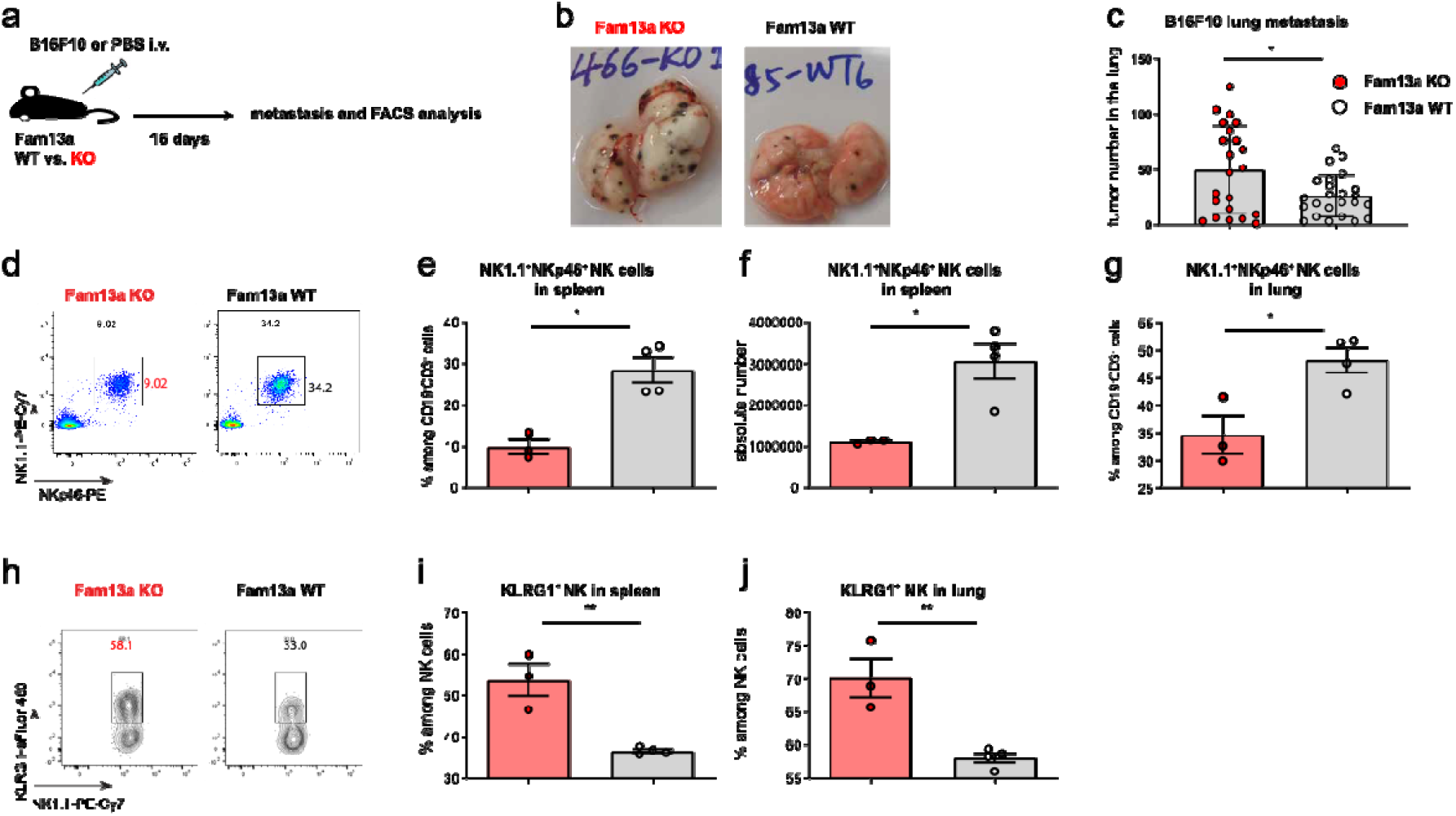
*Fam13a* deficiency aggravates lung metastasis induced by melanoma cells. **(a)** Schematic of the experimental setup for lung metastasis model induced by B16F10 melanoma cells. **(b)** Representative photographs of freshly isolated lungs after B16F10 cell inoculation. **(c)** Quantified metastatic foci of freshly isolated lungs after B16F10 cell inoculation (pooled KO, n=21; pooled WT, n=23). **(d)** Representative FACS plot of NKp46 and NK1.1 expression on *Fam13a* KO and WT CD3^−^CD19^−^ living singlet lymphocytes. **(e, g)** Percentage of NKp46^+^NK1.1^+^ NK cells out of CD3^−^CD19^−^ cells in spleen (**e**) or lung (**g**) of *Fam13a* KO and WT littermates. **(f)** Absolute number of NKp46^+^NK1.1^+^ NK cells out of CD3^−^CD19^−^ cells in spleen of *Fam13a* KO and WT littermates. **(h)** Representative FACS plot of KLRG1 and NK1.1 expression on *Fam13a* KO and WT NK cells in the spleen. **(i, j)** Percentage of KLRG1^+^ NK cells among total NK cells in spleen (**i**) or lung (**j**) (KO, n=3, WT, n=4). Data are mean ± s.d. The *p*-values were calculated by a two-tailed Student’s t-test. n.s. or unlabeled, not significant, \**p*<=0.05, \*\**p* <=0.01 and \*\*\**p* <=0.001.

Although NK cells are indispensable in suppressing B16F10-induced lung metastasis, we hitherto cannot exclude the potential role of other immune cells [30]. We therefore assayed lung-infiltrating T lymphocytes, CD4^+^ T cells and cytotoxic CD8^+^ T cells. We observed a higher percentage of CD4^+^ T cells and CD25^high^CD4^+^ Tregs **(S3a-b Fig)**, but not CD8 T cells (**S3c Fig**) in *Fam13a* KO compared to that in WT littermates. Furthermore, there was a higher frequency of effector memory (EM) but a lower percentage of naïve CD4^+^ T cells in *Fam13a* KO mice, indicating more CD4^+^ T cells were participating in the fight against melanoma, which could be either a driving factor or a secondary response of deteriorated lung metastasis (**S3d-e Fig**). This effect on naïve or memory compartment was not so obvious yet in infiltrated CD8^+^ T cells (**S3f Fig**). We also found there were much higher percentages of PD-1^+^ CD4^+^ T cells in the lung (**S3g-h Fig**), indicating more exhausted or activated CD4^+^ T cells in *Fam13a* KO mice. Again, the effect on PD-1 expression was not so significant in infiltrated CD8^+^ T cells of *Fam13a* KO mice (**S3i Fig**). These results indicate that, although a major role of *Fam13a*-deficient NK cells in the rejection of melanoma cells is likely, we cannot definitively exclude a potential contribution from other *Fam13a*-deficient immune cells, e.g., CD4^+^ T cells.

## Discussion

In this work, we showed that (i) *Fam13a* depletion upregulates the expression of the inhibitory and maturation marker, KLRG1 of NK cells, (ii) *Fam13a* is required for an optimal NK cell IFN-γ production, (iii) *in vivo*, the KO mice develop significantly more lung metastases after intravenous injection of a melanoma cell line, and (iv) the absence of *Fam13a* does not generate a phenotypic impact on B and T cell numbers and subset distribution, nor on Treg function (at least under homeostasis).

For *ex vivo* phenotyping analysis of *Fam13a*-KO mice, the most striking observation we found is the enhanced expression of KLRG1 in *Fam13a*-KO NK cells vs. WT NK cells. KLRG1 is an IR of the C-type lectin superfamily able to inhibit NK cell functions (cytotoxicity and cytokine production) upon recognition of its non-MHC class I ligands, which are E-, N- and R-cadherins [18, 22, 23]. The latter are adhesion molecules downregulated on cancer cells. In mice, KLRG1 is expressed on roughly a third of NK cells, but it increases upon infections [22, 23]. In *Fam13a*-KO mice, particularly in spleen, this value was almost doubled in most of the animals. Furthermore, KLRG1 is regarded as a maturation and terminal differentiation marker of NK cells. In line with this notion, we also observed modestly increased percentages of mature CD11b^+^CD27^−^ NK cells, which suggests that *Fam13a* has an inhibiting or regulatory effect on NK cell maturation or preventing to some extent premature entry into the senescent state.

Our *in vitro* NK cell functional studies showed that *Fam13a*-deficient NK cells displayed an impaired IFN-γ production following IL-2/IL-12/IL-15 or IL-18 cytokine cocktails stimulation. Furthermore, after an *in vivo* pre-activation of NK cells via the TLR3 agonist poly(I:C), *Fam13a*-KO NK cells produced less IFN-γ compared to WT NK cells. We also observed a mild effect of *Fam13a* depletion on NK degranulation, i.e., no significant difference in the percentages of CD107a expressing NK cells, except for a modest decrease in the MFI of CD107a^+^ NK cells. Since KLRG1 inhibits IFN-γ expression in NK cells at least in chronic viral infections [27–29], it is not surprising to observe impaired IFN-γ production in NK cells following various types of activation.

In our *in vivo* B16F10 melanoma models, we observed a strong reduction in the number of NK cells in spleen and lung of the KO mice compared to their WT counterparts, but a significantly higher frequency of KLRG1^+^ NK cells. This might suggest at first sight that more NK cells were inhibited following the induction of lung metastasis. Moreover, *Fam13a*-KO mice displayed a significantly higher number of lung metastases compared to the WT animals, which is in accordance with our *in vitro* findings, showing a hypofunctional state of KO NK cells. Since IFN-γ production by NK cells is critical to control B16F10-induced lung metastasis [31], the observed reduced capability of IFN-γ production in *Fam13a* KO NK cells, might at least partially contribute to more severe metastasis *in vivo*. In line with our observation about a higher expression of KLRG1 and the *in vitro* hypofunctional state of NK cells in *Fam13a* KO mice, KLRG1 neutralization antibody plus anti-PD-1 combinatory treatment vs. the anti-PD-1 therapy alone has achieved better survival rate and response to tumor volume in B16F10 melanoma models [32]. Furthermore, KLRG1^+^ NK cells possibly produced, as previously described, less IFN-γ than KLRG1^−^ NK cells in mice [19]. These data together can already well explain more severe metastasis observed in *Fam13a*-KO mice.

We also observed certain effects of *Fam13a* on filtrated T cells, especially on CD4^+^ T cells, in the induced lung metastasis experiments, so that further depletion experiments and cell-type specific knockout animal models might be required to figure out whether our observation is driven by NK cells alone or together with other cell types. In short, here we demonstrated a previously unrecognized critical role of *FAM13A* in regulating KLRG1 expression and IFN-γ production of NK cells using both *in vitro* and *in vivo* models. As *FAM13A* is strongly associated with the risk of several common lung diseases, our discovery paves the way to develop a novel potential target to mediate the functions of immune cells, especially NK cells in the fight against complex lung diseases, one of the major health threats globally. Encouragingly, the IFN-γ production in NK cells from patients with another *FAM13A*-strongly-linked lung disease, i.e., COPD, was reported to be impaired [33], indicating that our discovery might have translational potential.

## Materials and Methods

### Animals

*Fam13a* knock-out (KO) mice [Fam13a^tm2a (KOMP)Wtsi^] from the project *CSD70561* with C57BL/6N background were obtained from the Knockout Mouse Project (KOMP) Repository at UC Davis. The *Fam13a*^*-/-*^ (KO), *Fam13a*^*+/−*^ (HET) and *Fam13a*^*+/+*^ (WT) mice used in the experiments were age- and gender-matched littermates generated from *Fam13a*^*+/−*^ heterozygous breeding pairs. All mice were bred and maintained in our specific pathogen-free animal facilities. Both genders were used in our experiments depending on the availability. But we only employed one gender for each individual experiment.

### Ethics statement

All animal experimental protocols were performed following the approval of the Animal Welfare Structure (AWS) of the University of Luxembourg and the Luxembourg Institute of Health, and authorization from Luxembourg Ministry of Agriculture.

### Flow cytometry analysis of immunophenotype

Cell suspensions were obtained by the mechanical disruption of mouse spleen, peripheral lymph nodes, bone marrow and lung. Red blood cells were then lysed in 1x lysing buffer (555899, BD Biosciences). One million cells per sample were pre-incubated with purified anti-mouse CD16/CD32 antibody (Fc Block^™^, 553141, BD Biosciences). Live/dead cells were discriminated by staining cells with the LIVE/DEAD^®^ Fixable Near-IR Dead Cell Stain kit (L10119, Thermo Fisher Scientific) (dilution 1:500). Detailed antibody information was provide in Supplementary **Table S2**. Cell surface markers were stained by the combination of the following antibodies: anti-mouse CD3-BUV496 (612955, BD Biosciences) (dilution 1:100), anti-mouse CD3-BV421(562600, BD Biosciences) (dilution 1:100), anti-mouse CD19-BV510 (562956, BD Biosciences) (dilution 1:100), anti-mouse CD19-FITC (11-0193-82, eBioscience) (dilution 1:100), anti-mouse NK1.1-PE-Cy7 (108713, Biolegend) (dilution 1:50), anti-mouse NK1.1-BV421 (562921, BD Biosciences) (dilution 1:100), anti-mouse CD27-FITC (11-0271-82, eBioscience), anti-mouse CD11b-APC (553312, BD Biosciences) (dilution 1:100), anti-mouse CD11b-BUV395 (565976, BD Biosciences) (dilution 1:100), anti-mouse KLRG1-eFluor 450 (48-5893-82, eBioscience) (dilution 1:100), anti-mouse KLRG1-PerCP-Cy5.5 (138417, Biolegend) (dilution 1:100), anti-mouse CD69-PE (553237, BD Biosciences) (dilution 1:200), anti-mouse Ly49C/I-FITC (553276, BD Biosciences) (dilution 1:100), anti-mouse NKG2A-PE (142803, Biolegend) (dilution 1:100), anti-mouse CD69-BV605 (104529, Biolegend) (dilution 1:200), anti-mouse Ly49A-BUV395 (742370, BD Biosciences) (dilution 1:100), anti-mouse Ly49D-BV711 (742559, BD Biosciences) (dilution 1:100), anti-mouse 2B4-FITC (553305, BD Biosciences) (dilution 1:100), anti-mouse Ly49H-BUV395 (744266, BD Biosciences) (dilution 1:100), anti-mouse NKp46-PE (137603, Biolegend) (dilution 1:100), anti-mouse NKG2D-APC (130211, Biolegend) (dilution 1:100), anti-mouse CD4-FITC (11-0042-82, eBioscience) (dilution 1:200), anti-mouse CD8-BUV805 (564920, BD Biosciences) (dilution 1:200), anti-mouse CD25-APC (557192, BD Bioscience) (dilution 1:100), anti-mouse CD44-PE-Cy7 (560569, BD Biosciences) (dilution 1:200), anti-mouse CD62L-PerCP-Cy5.5 (560513, BD Biosciences) (dilution 1:200), anti-mouse PD-1-BV711 (744547, BD Biosciences) (dilution 1:200). Intracellular staining for Foxp3-APC (126403, Biolegend) (dilution 1:200), Ki-67-BV605 (652413, Biolegend) (dilution 1:200) or Ki-67-eFluor 450 (48-5698-82, eBioscience) (dilution 1:200) was performed by using the Foxp3 Staining Kit (00-5523-00, eBioscience). The choice of the combination of the antibodies was decided not only by the specific experimental purposes, spectrum compatibility, but also by our Fortessa laser settings. Samples were measured on a BD LSR Fortessa^™^ and data were analysed with FlowJo (v10, Tree Star).

### Primary murine NK cell *in vitro* expansion and cytokine activation

Five million of splenocytes per well in 12-well plates were cultured in 2.5 ml of complete DMEM medium (for details refer below) with 1000 U/ml of recombinant human (rh) IL-2 (202-IL-010, R&D Systems) at 37°C, in 5% CO_2_ incubators until day 5, when the cells were stimulated with 1000 U/ml of rhIL-2 and the combination of 10 ng/ml of recombinant murine (rm) IL-12 (210-12, PeproTech Inc) and 40 ng/ml of rm IL-15 (210-15, PeproTech Inc) or 100 ng/ml of rm IL-18 (B004-5, MBL) for one night. Before FACS staining, the cells were incubated with Golgistop (554724, BD Biosciences) (dilution 1:1500) and Golgiplug (555029, BD Biosciences) (dilution 1:1000) for 5 hrs. Cell surface markers were stained with anti-mouse CD3-BUV496 (612955, BD Biosciences) (dilution 1:100) and anti-mouse NK1.1-PE-CY7 (108713, Biolegend) (dilution 1:50). Dead cells were stained by using the LIVE/DEAD^®^ Fixable Near-IR Dead Cell Stain kit (dilution 1:500). For intracellular cytokine staining, cells were first fixed/permeabilized with Cytofix/Cytoperm buffer (554714, BD Biosciences) and then stained with anti-mouse IFN-γ-APC antibody (554413, BD Biosciences) (dilution 1:100) diluted in Perm/Wash buffer (554714, BD Biosciences). NK cell culture medium is the complete DMEM medium (41965039, Thermo Fisher Scientific) with 10% of Fetal Bovine Serum (FBS, 10500-064, Thermo Fisher Scientific), 100 U/ml Penicillin-Streptomycin (15070-063, Thermo Fisher Scientific), 10 mM HEPES (15630080, Thermo Fisher Scientific) and 55 µM β-mercaptoethanol (M7522, Sigma-Aldrich).

### NK cell degranulation assay

*Fam13a* KO and WT littermates were injected with 150 mg of toxin-free poly(I:C) (tlrl-pic, InvivoGen) in 100 µl PBS via intraperitoneal injection. Eighteen hours later, the mice were sacrificed and the spleens were harvested. Splenocytes (1E6) were cultured with 5E5 of YAC-1 target cells (2:1 ratio) in the presence of anti-mouse CD107a-BV421 antibody (564347, BD Biosciences) (1:100) for 1 hr. Golgiplug and Golgistop were then added for an additional 4-hr incubation. Cells were stained with anti-mouse CD3-BUV496 (612955, BD Biosciences) (dilution 1:100), anti-mouse NK1.1-PE-Cy7 (108713, Biolegend) (dilution 1:50) plus Live/Dead staining kit (dilution 1:500). Intracellular cytokine IFN-γ staining was performed as described above.

### Melanoma model for pulmonary metastasis

*Fam13a* KO and matched *Fam13a* WT littermate mice aged 8 to 10 weeks were used for lung metastasis study following intravenous injection of 2E5 of B16F10 melanoma cells in 100 µl of PBS. 16 days later, mice were sacrificed. The lung was perfused to remove the excessive blood by slowly injecting cold PBS through the right ventricle. Lung tumor nodules were counted. In addition, the spleens, pLNs and lungs were collected for FACS analysis. For cell isolation from the lung, the organs were cut into small pieces and the tissue digested in the lysis buffer (PBS containing 1.3 mg/ml collagenase Type II (234155, MERCK), 10% FBS, 50 U/ml benzonase endonuclease (101654, MERCK), 1mM MgCl_2_) in a 37 °C incubator for 1 h. Single cell suspensions for FACS staining were made by filtering of the cells through 40 µM cell strainers (734-2760, VWR).

### Real time PCR

Spleen and/or lung were collected from *Fam13a* KO, *Fam13a* Het and *Fam13a* WT mice. NK cells in the spleen were isolated and purified by using mouse NK cell isolation kit II (130-096-892, Miltenyi Biotec). The purity of NK cells was analyzed by FACS. RNA from lung and NK cells of spleen was isolated via using the RNeasy Mini Spin Kit (74104, Qiagen). Genomic DNA was then removed by on-column DNase digestion (79254, Qiagen). The SuperScript III Reverse Transcriptase Kit (18080-044, Invitrogen) was employed for cDNA synthesis. qPCR was performed with the LightCycler 480 SYBR Green I Master Mix (04707516001, Roche Applied Science) using the LightCycler 480 system as previously described [15, 34]. *Fam13a* mRNA RT-PCR primers are referred according to another *Fam13a* related work [35] (*fwd*: CCG CTG CGA AGC TCA CAG GAA GAT G; *rev*: TTG GTC TCC AGC GTT GCT GAC ATC A). The housekeeping gene for normalizing *Fam13a* mRNA expression was *Rps13* in lung and *18s* in splenic NK cells, respectively.

### Treg suppressive assay

Treg suppressive assay was performed in a way similar to our previous work [34]. To ease the comprehension, were here described the major steps again. Total T cells from spleen of *Fam13a* KO mice and WT littermates were isolated with CD90.2 microbeads (130-121-278, Miltenyi Biotec). The cells were then stained with the antibody mix containing LIVE/DEAD^®^ Fixable Near-IR (dilution 1:500), anti-mouse CD25-PE-Cy7 (552880, BD Biosciences) (dilution 1:200) and CD4-FITC (11-0042-82, eBioscience) (dilution 1:200) antibodies at 4 °C for 30 min. After staining, Tregs (CD4^+^CD25^high^) and conventional CD4^+^CD25^−^ T cells (Tconv) were sorted by BD FACSAria^™^ III sorter. Tconv cells from WT mice were labelled with a final concentration of 1 µM CellTrace^™^ CFSE (C34554, Life Technology). Splenocytes depleted of T cells were used as antigen-presenting cells (APCs)/feeder cells and were irradiated in RS2000 (Rad Source Technologies) with a total dose of 30 Gy within a period of 10 mins. 1×10^5^ Tconv cells were co-cultured with Treg cells at different ratios in the presence of 2×10^5^ irradiated APCs and 1 µg/ml soluble anti-CD3 antibody (554829, BD Biosciences). The cells were cultured in T-cell complete media which consists of RPMI 1640 medium supplemented with 50 U/ml penicillin, 10 mM HEPES, 10% heat-inactivated fetal bovine serum (FBS), 50 µg streptomycin, 2 mM GlutaMAX (35050061, Thermo Fisher Scientific), 1 mM sodium pyruvate (11360070, Thermo Fisher Scientific), 0.1 mM non-essential amino acids (M7145, Sigma Aldrich) and 50 µM beta-mercaptoethanol at 37°C, 5% CO_2_ incubator. The proliferation of Tconv cells labeled by CFSE was measured after three-day co-culturing using a BD LSR Fortessa^™^.

### Statistical analysis

The P-values were calculated by Graphpad Prism software with non-paired two-tailed Student t test as presented in the corresponding figure legends. The P-values under 0.05 were considered as statistically significant (*p < 0.05, **p < 0.01, ***p < 0.001; n.s, not significant). Data were presented as mean ± standard deviation (s.d.).

## Abbreviations

APC: Antigen-presenting cells
AR: Activating receptor
AWS: Animal welfare structure
CFSE: Carboxy fluorescein succinimidyl ester
CM: Central memory
COPD: Chronic obstructive lung disease
EM: Effector memory
eQTL: Expression quantitative trait loci
FACS: Flow cytometry
Fam13a: Family with sequence similarity 13, member A
FBS: Fetal bovine serum
GWAS: Genome-wide association study
HET: Heterozygous
IFN-γ: Interferon gamma
IL-2/-12/-15/-18: Interleukin 2/12/15/18
IR: Inhibitory receptor
KIR: Killer immunoglobulin-like receptors
KLRG1: Killer cell lectin-like receptor G1
KO: Knockout
MFI: Mean fluorescent intensity
MHC: Major histocompatibility complex
NK: Nature Killer
PBS: Phosphate-buffered saline
PCR: Polymerase chain reaction
pLN: peripheral lymph nodes
Tconv: Conventional CD4 T cells
Treg: CD4 regulatory T cells
WT: Wildtype

## Supporting Information

**Supplementary Table 1.** Frequency summary of unaffected immune subsets in homeostatic *Fam13a* KO vs WT mice. Data are provided as mean ± s.d.. The number of analyzed mice per group was also provided.

|  | Fam13a KO | Fam13a WT |
| --- | --- | --- |
| **cell type (in Spleen)** | **% among the parent gate (% , mean± standard deviation)** |  |
| CD3<sup>+</sup> T cells (KO=12,WT=11) | 32.28±3.38 | 29.29±8.73 |
| CD19<sup>+</sup> B cell (KO=12,WT=11) | 48.6±11.61 | 55.34±11.67 |
| CD4<sup>+</sup> T cells (KO,WT N=5) | 12.99±2.29 | 12.23±2.11 |
| CD44<sup>low</sup>CD62L<sup>high</sup> naïve CD4 (KO,WT N=5) | 46.35±15.28 | 53.75±3.31 |
| CD44<sup>high</sup>CD62L<sup>low</sup> EM CD4 (KO,WT N=5) | 34.45±14.3 | 28.80±3.00 |
| CD69<sup>+</sup>CD4<sup>+</sup> T cells (KO,WT N=5) | 9.73±1.35 | 11.91±0.96 |
| KI-67<sup>+</sup>CD4<sup>+</sup> T cells (KO,WT N=5) | 13.85±4.47 | 12.36±1.20 |
| PD-1<sup>+</sup>CD4<sup>+</sup> T cells (KO,WT N=5) | 19.47±6.47 | 18.16±3.49 |
| FOXP3<sup>+</sup>CD4<sup>+</sup> Tregs (KO,WT N=5) | 19.03±5.40 | 15.66±1.87 |
| CD8<sup>+</sup> T cells (KO,WT N=5) | 9.89±1.75 | 8.70±1.77 |
| naïve CD8<sup>+</sup> T cells (KO,WT N=5) | 53.08±16.35 | 64.14±1.62 |
| EM CD8<sup>+</sup> T cells (KO,WT N=5) | 3.85±1.95 | 2.78±0.79 |
| CD44<sup>high</sup>CD62L<sup>high</sup> CM CD8<sup>+</sup> T cells (KO,WT N=5) | 37.87±16.42 | 27.53±0.95 |
| CD69<sup>+</sup>CD8<sup>+</sup> T cells (KO,WT N=5) | 1.81±0.40 | 2.07±0.35 |
| Ki-67<sup>+</sup>CD8<sup>+</sup> T cells (KO,WT N=5) | 7.23±1.60 | 6.94±0.71 |
| PD-1<sup>+</sup>CD8<sup>+</sup> T cells (KO,WT N=5) | 2.75±0.76 | 2.29±0.20 |

**Supplementary Table 2.**
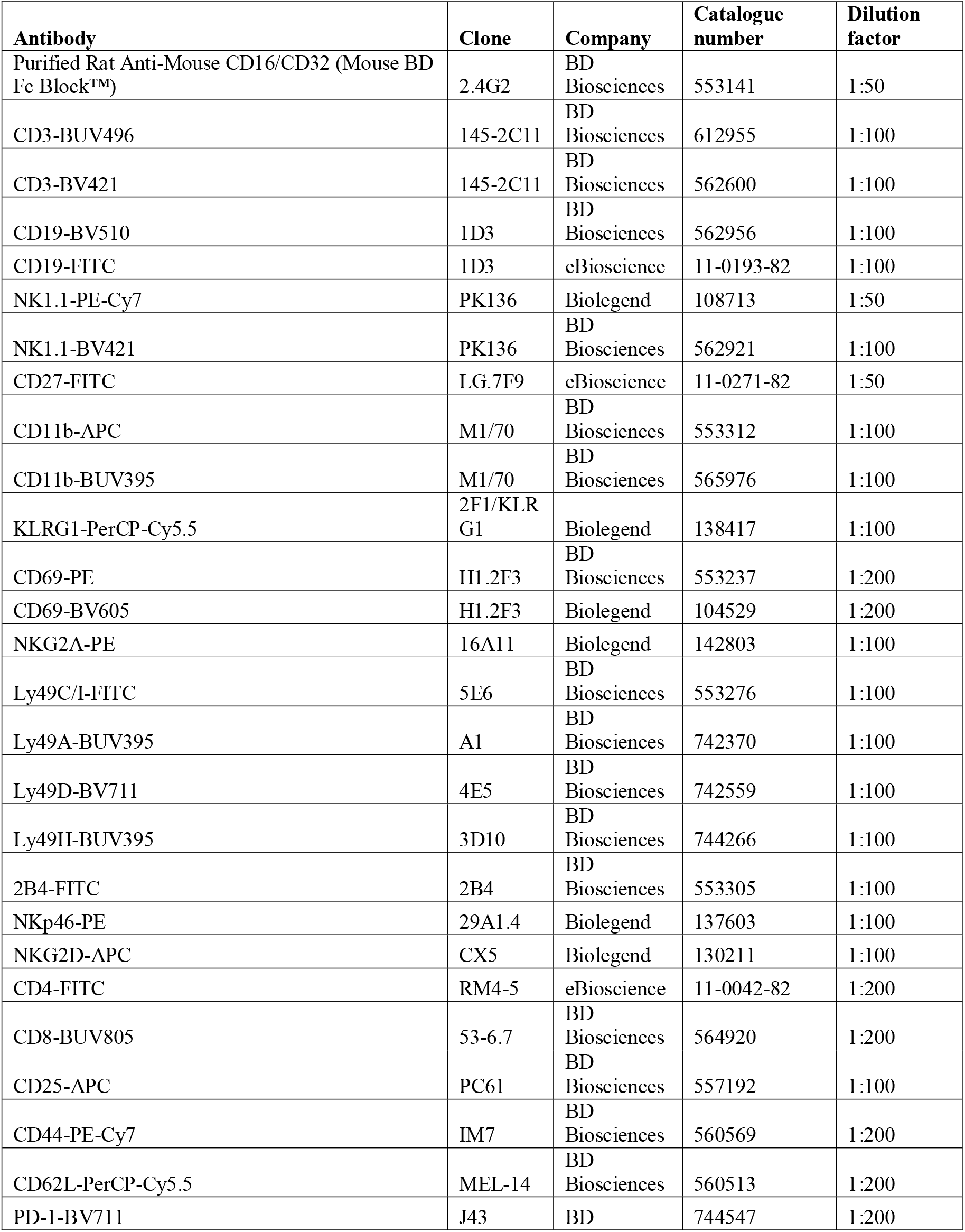
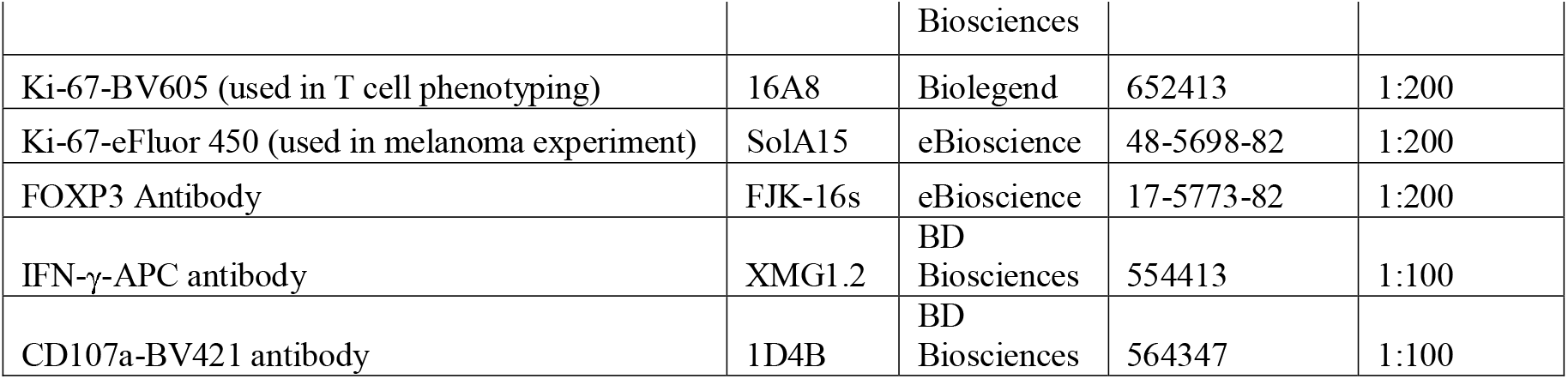
List of antibodies with the information required for flow cytometry analysis.

**S1 Fig.**
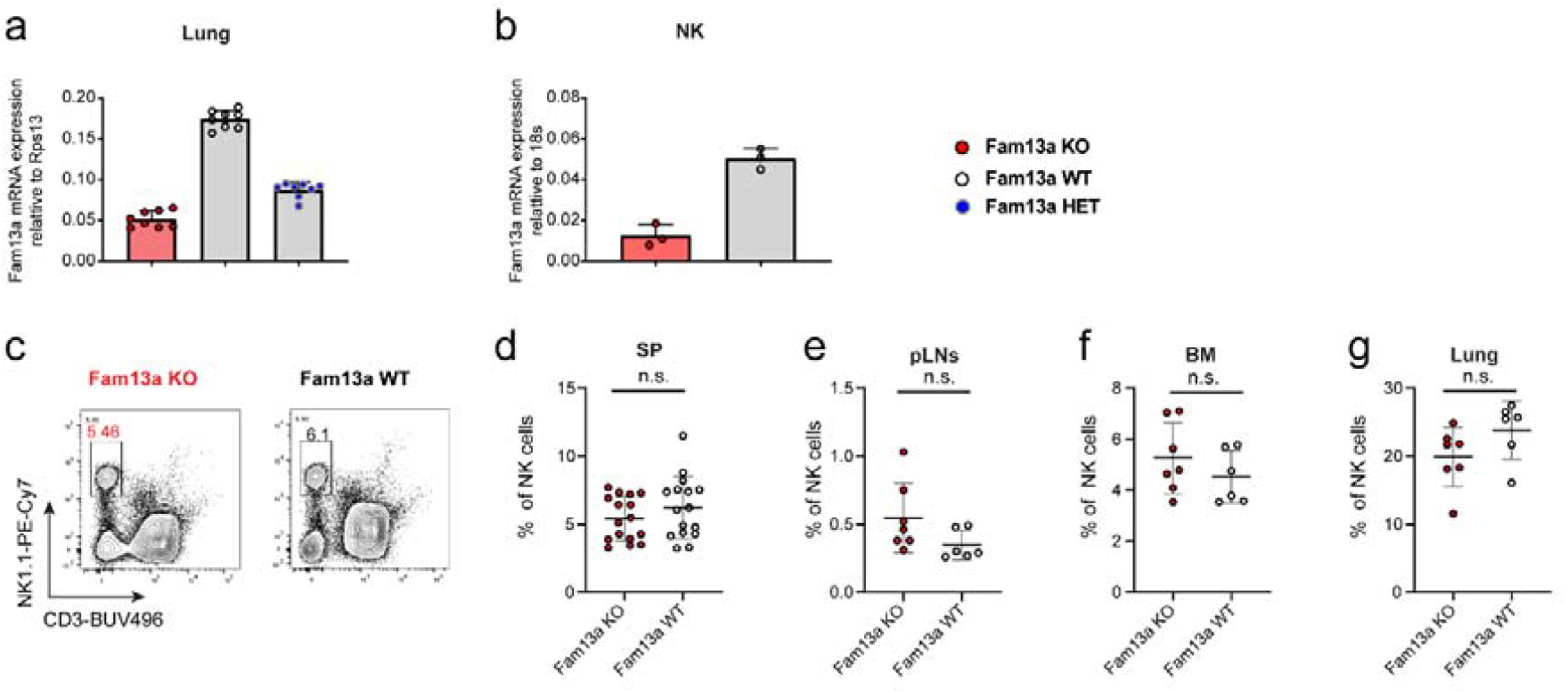
Frequency of NK cells in the different organs and tissues of *Fam13a* KO vs. WT mice. **(a-b)** Relative *Fam13a* mRNA expression in lung tissue (**a**) or fresh NK cells (**b**) isolated from *Fam13a* KO or WT littermates or Fam13a heterozygous (HET) mice as quantified by real-time PCR. **(c)** Representative flow-cytometry (FACS) plots of CD3^−^ NK1.1^+^ NK cells among CD19^−^ cells in spleen of *Fam13a* KO and WT littermates. **(d-g)** Percentages of NK cells in spleen (**d**, KO, n=16; WT, n=16), peripheral lymph nodes (pLNs) (**e**, KO, n=7; WT, n=6), bone marrow (BM) (**f**, KO, n=7; WT, n=6) and lung (**g**, KO, n=7; WT, n=6) of *Fam13a* KO and WT littermates. Results are representative of three (**c-g**) and two (**a-b**) independent experiments. Data are mean ± s.d. The *p*-values were calculated by a two-tailed Student’s t-test. n.s. or unlabeled, not significant, \**p*<=0.05, \*\**p* <=0.01 and \*\*\**p* <=0.001.

**S2 Fig.**
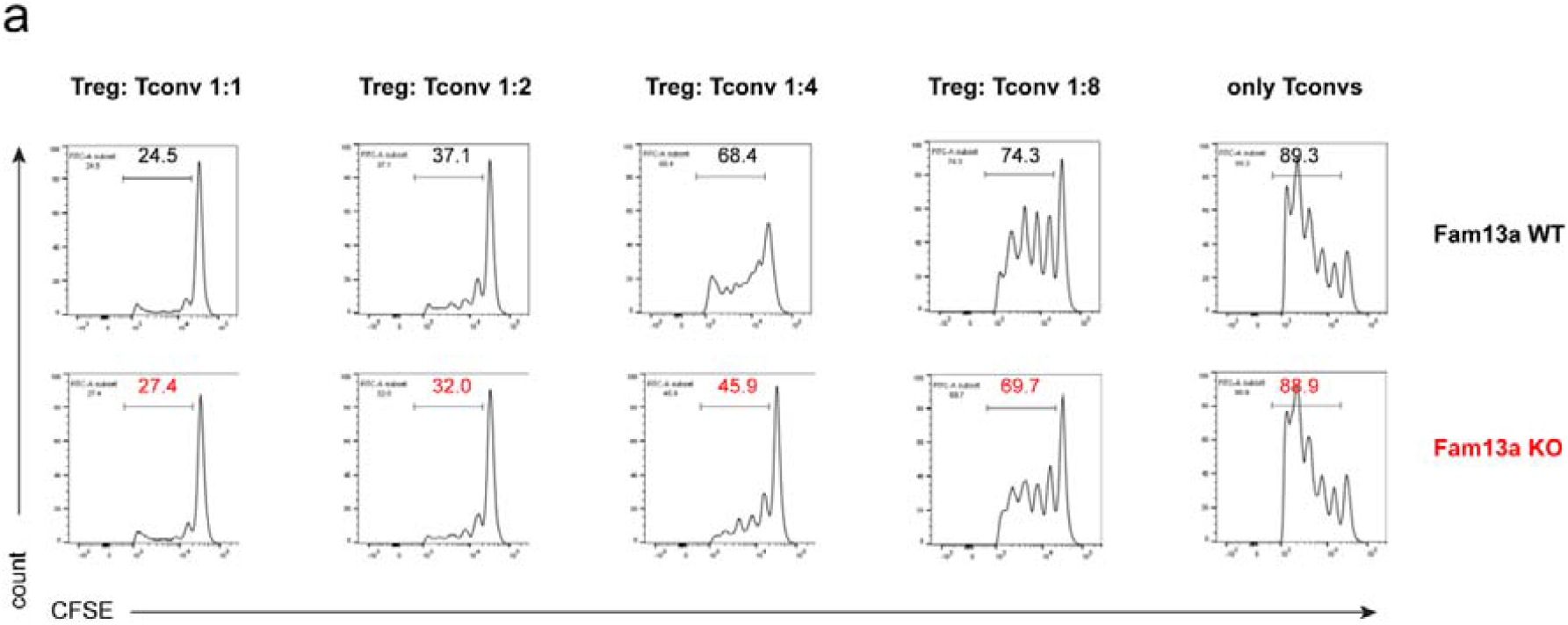
Treg suppressive assay. **(a)** *In-vitro* suppression assay of *Fam13a* KO or WT Tregs in co-culture with CFSE-labelled Tconv cells at different ratios and irradiated feeder cells in the presence of anti-CD3 antibody for 3 days. Enlarged number in each histogram represents percentage of dividing cells from the total living Tconv population. Results represent four independent experiments.

**S3 Fig.**
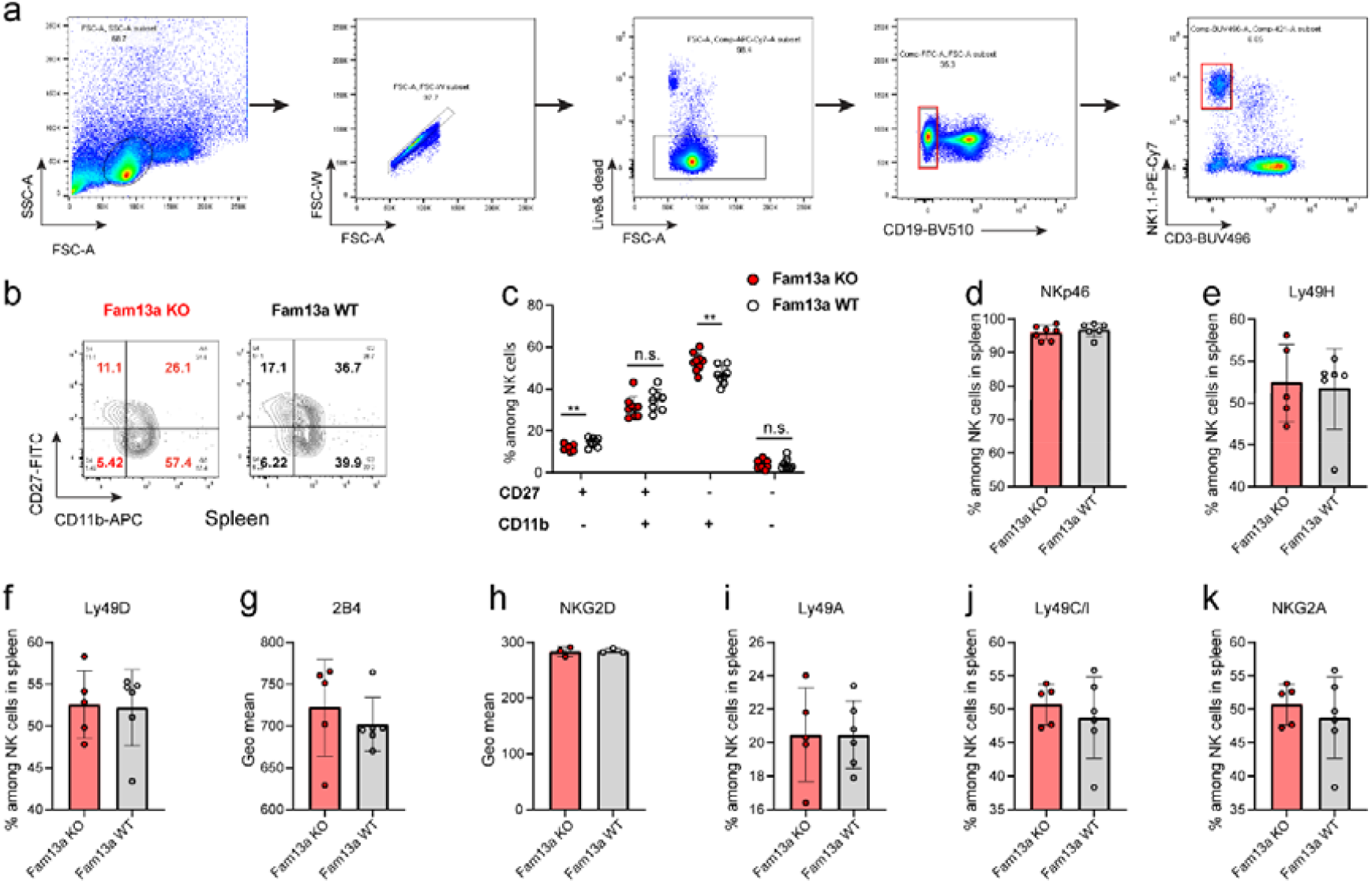
Extended NK cell immunophenotyping analysis of *Fam13a* KO vs. WT mice. **(a)** Representative FACS gating strategy of NK cells from splenocyte singlets stained with live and dead staining dye, anti-CD19, anti-CD3 and anti-NK1.1antibodies. **(b)** Representative FACS plots of CD27 and CD11b expression gated on CD19^−^CD3^−^ NK1.1^+^cells in spleen. **(c)** Percentages of four developmental stages of NK cells, CD27^−^CD11b^−^, CD27^+^CD11b^−^, CD27^+^CD11b^+^, CD27^−^CD11b^+^ NK cells in spleen of *Fam13a* KO and WT littermates (KO, n=9; WT, n=8). **(d-f)** Percentages of NK cells expressing activating receptors NKp46 (**d**), Ly49H (**e**) and Ly49D (**f**) among *Fam13a* KO and WT littermates (KO, n=5; WT, n=6). **(g-h)** Geometric mean of 2B4 (**g**) and NKG2D (**h**) expression on *Fam13a* KO and WT NK cells. **(i-k)** Percentages of NK cells expression inhibitory receptors Ly49A (**i**), Ly49C/I (**j**) and NKG2A (**k**) among *Fam13a* KO and WT littermates (KO, n=5; WT, n=6). Results represent four (**c**) and three (**d-k**) independent experiments. Data are mean± s.d. The p-values were determined by a two-tailed Student’s t-test. n.s. or unlabeled, not significant, *p<=0.05, **p<=0.01 and ***p<=0.001.

**S4 Fig.**
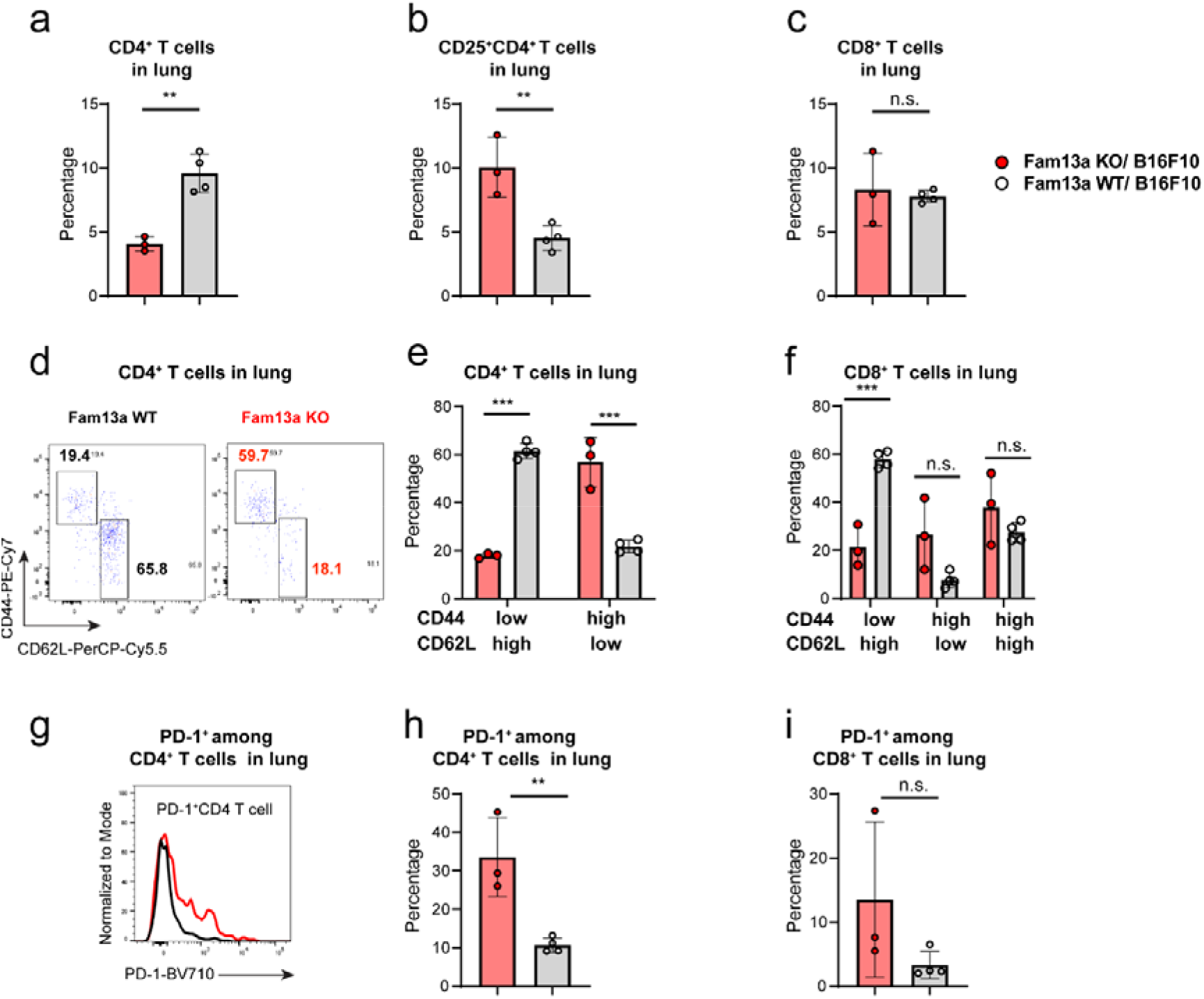
Extended analysis of immune cells in the B16F10-melanoma lung metastasis model. **(a-c)** Frequency of CD4^+^ T cells **(a)**, CD25^hi^CD4^+^ Tregs (**b**) and CD8^+^ T cells (**c**) in the lung of *Fam13a* KO and WT littermates. **(d)** Representative FACS plot of naïve (CD62L^hi^ CD44^low^) and effector memory (EM) (CD62L^low^ CD44^hi^ CD4^+^ T cells in the lung of *Fam13a* KO and WT littermates. **(e)** Percentage of naïve and EM among total CD4^+^ T cells in the lung of *Fam13a* KO and WT littermates. **(f)** Percentage of naïve, EM and CM among total CD8^+^ T cells in the lung of *Fam13a* KO and WT littermates. **(g)** Representative histogram overlays of PD-1 expression in CD4^+^ T cells in the lung of *Fam13a* KO and WT littermates. **(h-i)** Percentage of PD-1^+^ CD4^+^ T cells (**h**) and PD-1^+^CD8^+^ T cells (**i**) in the lung of *Fam13a* KO and WT littermates. Results represent two independent experiments. Data are mean ± s.d. The *p*-values were calculated by a two-tailed Student’s t-test. n.s. or unlabeled, not significant, \**p*<=0.05, \*\**p* <=0.01 and \*\*\**p* <=0.001.

## Acknowledgements

F.Q.H. and the group was partially supported by Luxembourg National Research Fund (FNR) CORE programme grant (CORE/14/BM/8231540/GeDES), FNR AFR-RIKEN bilateral programme (TregBAR, F.Q.H. and M.O.), PRIDE programme grants (PRIDE/11012546/NEXTIMMUNE and PRIDE/10907093/CRITICS). The students N.Z., C.C. and N.D.P. were supported through FNR AFR and DTU PhD programmes (PHD-2015-1/9989160, PRIDE/10907093/CRITICS and PRIDE/11012546/NEXTIMMUNE, respectively). The work was also partially supported through intramural funding of LIH and LCSB through Ministry of Higher Education and Research (MESR) of Luxembourg. The funders had no role in study design, data collection and analysis, decision to publish, or preparation of the manuscript.

## Author contributions

**Conceptualization:** Jacques Zimmer, Feng Q. Hefeng

**Funding acquisition:** Markus Ollert, Rudi Balling, Jacques Zimmer, Feng Q. Hefeng

**Investigation:** Ni Zeng, Maud Theresine, Christophe Capelle, Neha D. Patil, Cécile Masquelier, Caroline Davril, Alexandre Baron, Djalil Coowar, Xavier Dervillez, Aurélie Poli, Cathy Leonard, Jacques Zimmer, Feng Q. Hefeng

**Methodology:** Ni Zeng, Maud Theresine, Christophe Capelle, Cécile Masquelier, Caroline Davril, Alexandre Baron, Jacques Zimmer, Feng Q. Hefeng

**Project administration:** Feng Q. Hefeng

**Resources:** Markus Ollert, Rudi Balling, Jacques Zimmer, Feng Q. Hefeng

**Supervision:** Markus Ollert, Rudi Balling, Jacques Zimmer, Feng Q. Hefeng

**Visualization:** Ni Zeng, Jacques Zimmer, Feng Q. Hefeng

**Writing – original draft:** Ni Zeng, Jacques Zimmer, Feng Q. Hefeng

**Writing – review & editing:** Ni Zeng, Maud Theresine, Christophe Capelle, Neha D. Patil, Cécile Masquelier, Caroline Davril, Alexandre Baron, Djalil Coowar, Xavier Dervillez, Aurélie Poli, Cathy Leonard, Rudi Balling, Markus Ollert, Jacques Zimmer, Feng Q. Hefeng

## References

1. Yus Y, Mao L, Lu X, Yuan W, Chen Y, Jiang L, et al. Functional Variant in 3′ UTR of FAM13A Is Potentially Associated with Susceptibility and Survival of Lung Squamous Carcinoma. DNA and cell biology. 2019;38(11):1269–77.

2. Ziółkowska-Suchanek I, Mosor M, Podralska M, Iżykowska K, Gabryel P, Dyszkiewicz W, et al. FAM13A as a Novel Hypoxia-Induced Gene in Non-Small Cell Lung Cancer. Journal of Cancer. 2017;8(19):3933.

3. Castaldi PJ, Guo F, Qiao D, Du F, Naing ZZC, Li Y, et al. Identification of Functional Variants in the FAM13A Chronic Obstructive Pulmonary Disease Genome-Wide Association Study Locus by Massively Parallel Reporter Assays. American journal of respiratory and critical care medicine. 2019;199(1):52–61. Epub 2018/08/07. doi: 10.1164/rccm.201802-0337OC. PubMed PMID: 30079747; PubMed Central PMCID: PMCPMC6353020.

4. Hirano C, Ohshimo S, Horimasu Y, Iwamoto H, Fujitaka K, Hamada H, et al. FAM13A polymorphism as a prognostic factor in patients with idiopathic pulmonary fibrosis. Respiratory medicine. 2017;123:105–9. Epub 2017/02/01. doi: 10.1016/j.rmed.2016.12.007. PubMed PMID: 28137485.

5. Zhang Y, Qiu J, Zhang P, Zhang J, Jiang M, Ma Z. Genetic variants in FAM13A and IREB2 are associated with the susceptibility to COPD in a Chinese rural population: a casecontrol study. International journal of chronic obstructive pulmonary disease. 2018;13:1735-45. Epub 2018/06/07. doi: 10.2147/copd.S162241. PubMed PMID: 29872291; PubMed Central PMCID: PMCPMC5973397.

6. Wang B, Liang B, Yang J, Xiao J, Ma C, Xu S, et al. Association of FAM13A polymorphisms with COPD and COPD-related phenotypes in Han Chinese. Clinical biochemistry. 2013;46(16-17):1683–8. Epub 2013/07/31. doi: 10.1016/j.clinbiochem.2013.07.013. PubMed PMID: 23891779.

7. Ziółkowska-Suchanek I, Mosor M, Gabryel P, Grabicki M, Żurawek M, Fichna M, et al. Susceptibility loci in lung cancer and COPD: association of IREB2 and FAM13A with pulmonary diseases. Scientific reports. 2015;5:13502.

8. Young RP, Hopkins RJ, Hay BA, Whittington CF, Epton MJ, Gamble GD. FAM13A locus in COPD is independently associated with lung cancer–evidence of a molecular genetic link between COPD and lung cancer. The application of clinical genetics. 2011;4:1.

9. Kim S, Kim H, Cho N, Lee SK, Han BG, Sull JW, et al. Identification of FAM13A gene associated with the ratio of FEV1 to FVC in Korean population by genome-wide association studies including gene-environment interactions. Journal of human genetics. 2015;60(3):139–45. Epub 2015/01/23. doi: 10.1038/jhg.2014.118. PubMed PMID: 25608829.

10. Cho MH, Boutaoui N, Klanderman BJ, Sylvia JS, Ziniti JP, Hersh CP, et al. Variants in FAM13A are associated with chronic obstructive pulmonary disease. Nature genetics. 2010;42(3):200–2. Epub 2010/02/23. doi: 10.1038/ng.535. PubMed PMID: 20173748; PubMed Central PMCID: PMCPMC2828499.

11. van der Plaat DA, de Jong K, Lahousse L, Faiz A, Vonk JM, van Diemen CC, et al. Genome-wide association study on the FEV(1)/FVC ratio in never-smokers identifies HHIP and FAM13A. The Journal of allergy and clinical immunology. 2017;139(2):533–40. Epub 2016/09/11. doi: 10.1016/j.jaci.2016.06.062. PubMed PMID: 27612410.

12. Guo Y, Lin H, Gao K, Xu H, Deng X, Zhang Q, et al. Genetic analysis of IREB2, FAM13A and XRCC5 variants in Chinese Han patients with chronic obstructive pulmonary disease. Biochemical and biophysical research communications. 2011;415(2):284–7.

13. Jiang Z, Lao T, Qiu W, Polverino F, Gupta K, Guo F, et al. A chronic obstructive pulmonary disease susceptibility gene, FAM13A, regulates protein stability of β-catenin. American journal of respiratory and critical care medicine. 2016;194(2):185–97.

14. Eisenhut F, Heim L, Trump S, Mittler S, Sopel N, Andreev K, et al. FAM13A is associated with non-small cell lung cancer (NSCLC) progression and controls tumor cell proliferation and survival. Oncoimmunology. 2017;6(1):e1256526. Epub 2017/02/16. doi: 10.1080/2162402x.2016.1256526. PubMed PMID: 28197372; PubMed Central PMCID: PMCPMC5283630.

15. He F, Chen H, Probst-Kepper M, Geffers R, Eifes S, Del Sol A, et al. PLAU inferred from a correlation network is critical for suppressor function of regulatory T cells. Mol Syst Biol. 2012;8:624. Epub 2012/11/22. doi: 10.1038/msb.2012.56. PubMed PMID: 23169000; PubMed Central PMCID: PMCPMC3531908.

16. Schmiedel BJ, Singh D, Madrigal A, Valdovino-Gonzalez AG, White BM, Zapardiel-Gonzalo J, et al. Impact of Genetic Polymorphisms on Human Immune Cell Gene Expression. Cell. 2018;175(6):1701–15 e16. Epub 2018/11/20. doi: 10.1016/j.cell.2018.10.022. PubMed PMID: 30449622; PubMed Central PMCID: PMCPMC6289654.

17. Yao MY, Zhang WH, Ma WT, Liu QH, Xing LH, Zhao GF. microRNA-328 in exosomes derived from M2 macrophages exerts a promotive effect on the progression of pulmonary fibrosis via FAM13A in a rat model. Experimental & molecular medicine. 2019;51(6):1–16. Epub 2019/06/06. doi: 10.1038/s12276-019-0255-x. PubMed PMID: 31164635; PubMed Central PMCID: PMCPMC6547742.

18. Ito M, Maruyama T, Saito N, Koganei S, Yamamoto K, Matsumoto N. Killer cell lectinlike receptor G1 binds three members of the classical cadherin family to inhibit NK cell cytotoxicity. J Exp Med. 2006;203(2):289–95. Epub 2006/02/08. doi: 10.1084/jem.20051986. PubMed PMID: 16461340; PubMed Central PMCID: PMCPMC2118217.

19. Huntington ND, Tabarias H, Fairfax K, Brady J, Hayakawa Y, Degli-Esposti MA, et al. NK cell maturation and peripheral homeostasis is associated with KLRG1 up-regulation. the Journal of Immunology. 2007;178(8):4764–70.

20. Chiossone L, Chaix J, Fuseri N, Roth C, Vivier E, Walzer T. Maturation of mouse NK cells is a 4-stage developmental program. Blood. 2009;113(22):5488–96. Epub 2009/02/24. doi: 10.1182/blood-2008-10-187179. PubMed PMID: 19234143.

21. Elpek KG, Rubinstein MP, Bellemare-Pelletier A, Goldrath AW, Turley SJ. Mature natural killer cells with phenotypic and functional alterations accumulate upon sustained stimulation with IL-15/IL-15Ralpha complexes. Proc Natl Acad Sci U S A. 2010;107(50):21647–52. Epub 2010/11/22. doi: 10.1073/pnas.1012128107. PubMed PMID: 21098276.

22. Banh C, Fugere C, Brossay L. Immunoregulatory functions of KLRG1 cadherin interactions are dependent on forward and reverse signaling. Blood. 2009;114(26):5299–306. Epub 2009/10/27. doi: 10.1182/blood-2009-06-228353. PubMed PMID: 19855082; PubMed Central PMCID: PMCPMC2796135.

23. Li Y, Hofmann M, Wang Q, Teng L, Chlewicki LK, Pircher H, et al. Structure of natural killer cell receptor KLRG1 bound to E-cadherin reveals basis for MHC-independent missing self recognition. Immunity. 2009;31(1):35–46. Epub 2009/07/17. doi: 10.1016/j.immuni.2009.04.019. PubMed PMID: 19604491; PubMed Central PMCID: PMCPMC3030123.

24. Alter G, Malenfant JM, Altfeld M. CD107a as a functional marker for the identification of natural killer cell activity. Journal of immunological methods. 2004;294(1-2):15–22.

25. Aktas E, Kucuksezer UC, Bilgic S, Erten G, Deniz G. Relationship between CD107a expression and cytotoxic activity. Cellular immunology. 2009;254(2):149–54.

26. López-Soto A, Gonzalez S, Smyth MJ, Galluzzi L. Control of Metastasis by NK Cells. Cancer Cell. 2017;32(2):135–54. doi: https://doi.org/10.1016/j.ccell.2017.06.009.

27. Alvarez M, Simonetta F, Baker J, Pierini A, Wenokur AS, Morrison AR, et al. Regulation of murine NK cell exhaustion through the activation of the DNA damage repair pathway. JCI insight. 2019;4(14).

28. Müller-Durovic B, Lanna A, Covre LP, Mills RS, Henson SM, Akbar AN. Killer Cell Lectin-like Receptor G1 Inhibits NK Cell Function through Activation of Adenosine 5′Monophosphate–Activated Protein Kinase. The Journal of Immunology. 2016;197(7):2891–9.

29. Wang JM, Cheng YQ, Shi L, Ying RS, Wu XY, Li GY, et al. KLRG1 negatively regulates natural killer cell functions through the Akt pathway in individuals with chronic hepatitis C virus infection. Journal of virology. 2013;87(21):11626–36.

30. Souza-Fonseca-Guimaraes F, Cursons J, Huntington ND. The Emergence of Natural Killer Cells as a Major Target in Cancer Immunotherapy. Trends Immunol. 2019;40(2):142–58. Epub 2019/01/15. doi: 10.1016/j.it.2018.12.003. PubMed PMID: 30639050.

31. Takeda K, Nakayama M, Sakaki M, Hayakawa Y, Imawari M, Ogasawara K, et al. IFN-γ production by lung NK cells is critical for the natural resistance to pulmonary metastasis of B16 melanoma in mice. Journal of Leukocyte Biology. 2011;90(4):777–85. doi: 10.1189/jlb.0411208.

32. Greenberg SA, Kong SW, Thompson E, Gulla SV. Co-inhibitory T cell receptor KLRG1: human cancer expression and efficacy of neutralization in murine cancer models. Oncotarget. 2019;10(14):1399–406. Epub 2019/03/13. doi: 10.18632/oncotarget.26659. PubMed PMID: 30858925; PubMed Central PMCID: PMCPMC6402715.

33. Tang Y, Li X, Wang M, Zou Q, Zhao S, Sun B, et al. Increased numbers of NK cells, NKT-like cells, and NK inhibitory receptors in peripheral blood of patients with chronic obstructive pulmonary disease. Clin Dev Immunol. 2013;2013:721782. Epub 2013/09/27. doi: 10.1155/2013/721782. PubMed PMID: 24069043; PubMed Central PMCID: PMCPMC3773417.

34. Danileviciute E, Zeng N, Capelle C, Paczia N, Gillespie MA, Kurniawan H, et al. PARK7/DJ-1 promotes pyruvate dehydrogenase activity and maintains Treg homeostasis. bioRxiv. 2019:2019.12.20.884809. doi: 10.1101/2019.12.20.884809.

35. Wardhana DA, Ikeda K, Barinda AJ, Nugroho DB, Qurania KR, Yagi K, et al. Family with sequence similarity 13, member A modulates adipocyte insulin signaling and preserves systemic metabolic homeostasis. Proceedings of the National Academy of Sciences. 2018;115(7):1529–34.

